# The population frequency of human mitochondrial DNA variants is highly dependent upon mutational bias

**DOI:** 10.1101/2021.05.12.443844

**Authors:** Cory D. Dunn

**Affiliations:** Institute of Biotechnology, University of Helsinki, Helsinki, 00014, Finland

## Abstract

Next-generation sequencing can quickly reveal genetic variation potentially linked to heritable disease. As databases encompassing human variation continue to expand, rare variants have been of high interest, since the frequency of a variant is expected to be low if the genetic change leads to a loss of fitness or fecundity. However, the use of variant frequency when seeking genomic changes linked to disease remains very challenging. Here, we explore the role of selection in controlling human variant frequency using the HelixMT database, which encompasses hundreds of thousands of mitochondrial DNA (mtDNA) samples. We find that a substantial number of synonymous substitutions, which have no effect on protein sequence, were never encountered in this large study, while many other synonymous changes are found at very low frequencies. Further analyses of human and mammalian mtDNA datasets indicate that the population frequency of synonymous variants is predominantly determined by mutational biases rather than by strong selection acting upon nucleotide choice. Our work has important implications that extend to the interpretation of variant frequency for non-synonymous substitutions.

## Introduction

In this era of genomic medicine, next-generation sequence data obtained from patients, families, and populations are used to reveal and predict which genes and variants may be linked to disease (Claussnitzer et al., 2020; Shendure et al., 2019). Researchers have often embraced rare variants when seeking genomic changes in protein-coding sequences that may be pathogenic, as a reduction in variant frequency is an expected outcome of selection (Bomba et al., 2017; Gibson, 2012; Sazonovs and Barrett, 2018; Zuk et al., 2014). However, while triage of rare variants has led to some success in illuminating genes linked to heritable disease (Fuchsberger et al., 2016; Genovese et al., 2016; Lencz et al., 2021; Luo et al., 2017), the interpretation and utilization of rare genomic changes remains very challenging (Macklin et al., 2018; Manrai et al., 2016; Uricchio et al., 2016).

Human mitochondrial DNA (mtDNA) encodes proteins and RNAs required for the process of oxidative phosphorylation, and mitochondrial mutations are linked to a number of metabolic diseases (Gorman et al., 2016; Thompson et al., 2020). Recently, the mtDNAs of nearly 200,000 individuals were sequenced in order to produce the HelixMT database (HelixMTdb), a large catalog of human mtDNA variation (Bolze et al., 2019). Here, we find that many synonymous nucleotide substitutions were never detected within this quite substantial survey of human mtDNA. Subsequent study of more than one thousand mammalian mtDNAs suggested that selection on synonymous base substitution in mitochondrial protein-coding genes is minimal and unlikely to explain the absence or rarity of many synonymous changes within the human population. Rather, the mutational propensities of mtDNA (Kumar, 1996; Reyes et al., 1998) are more likely to have a dominant influence upon variant frequency. My findings have general implications for the interpretation of variant frequencies when studying heritable disease.

## Results

During an exploration of selective pressures that may act upon mitochondria-encoded polypeptides, I simulated every possible nucleotide substitution from the human reference mtDNA sequence within all protein-coding sequences, then cross-referenced these nucleotide and potential amino acid changes with the nucleotide substitutions tabulated in HelixMTdb. Consistent with selection against the vast majority of amino acid changes, non-synonymous substitutions were depleted to a far greater extent than synonymous substitutions when considering either the number of samples harboring variants of each type (Fig. 1a) or whether the nucleotide substitution was encountered at all during compilation of the HelixMTdb (Fig. 1b).

**Fig. 1:**
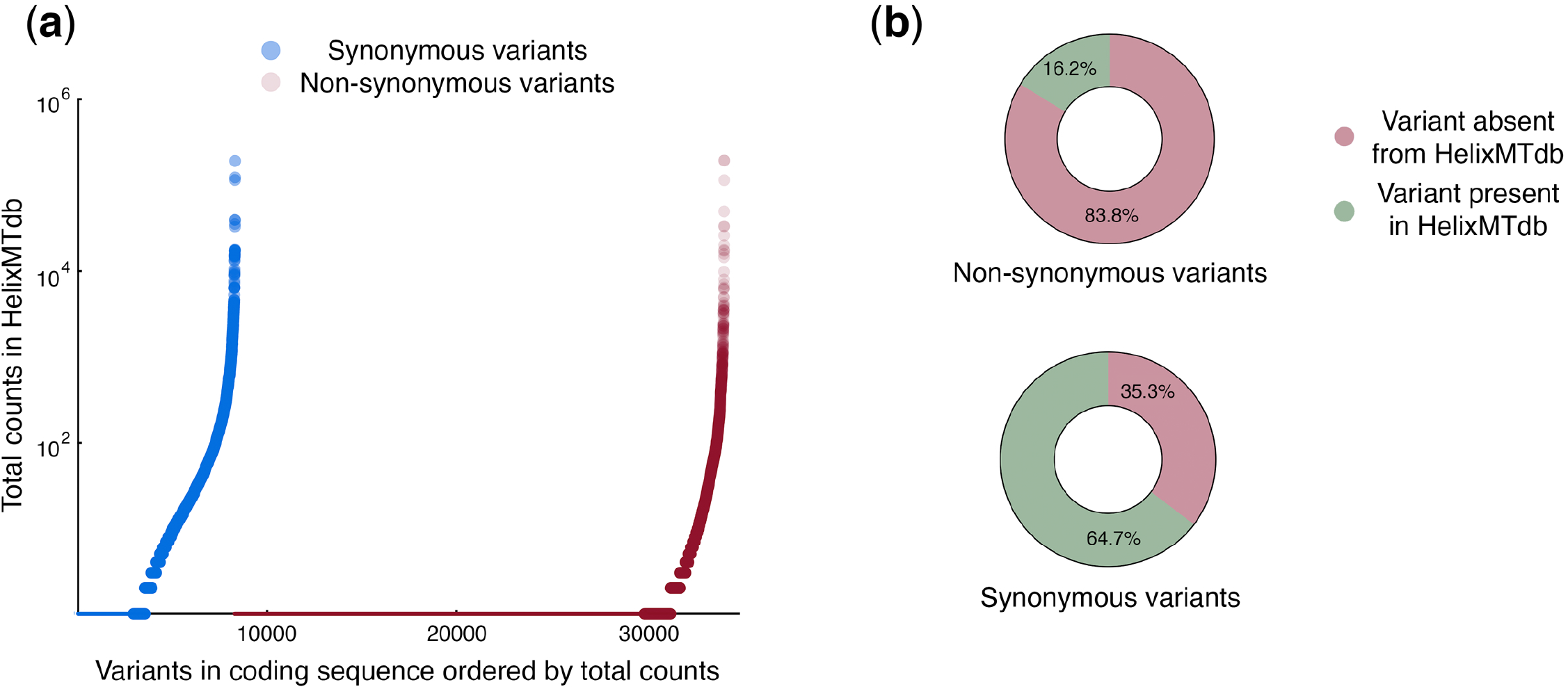
Many synonymous variants are never encountered within the HelixMT database. **(a)** Non-synonymous and synonymous variants differ in their population frequency. The sample count for variants within coding regions that are reachable by a single substitution from the human mtDNA reference sequence were obtained from HelixMTdb and plotted (Kolmogorov-Smirnov approximate P-value, <0.0001). Non-synonymous and synonymous substitutions never encountered are reflected as smaller dots along the x-axis at the zero position on the y-axis. **(b)** While the great majority of non-synonymous substitutions were never encountered during the HelixMTdb study, a substantial fraction of synonymous substitutions are also apparently lacking from the human population.

Since synonymous changes would not be expected to change the structure or function of mitochondria-encoded proteins, the number of changes (2925, or ∼35% of potential synonymous substitutions) for which a <u>s</u>ynonymous <u>s</u>ubstitution was <u>n</u>ever <u>e</u>ncountered (abbreviated here as ‘SSNEs’) during the HelixMTdb investigation was considered noteworthy. While HelixMTdb contained samples from almost all human haplogroups (Bolze et al., 2019), and while closely related individuals were removed from the study, more than 90% of samples were classified within the ‘N’ macro-haplogroup, a lineage associated with the exit of modern humans from Africa (Ingman et al., 2000; Maca-Meyer et al., 2001). Consequently, it was possible that SSNE abundance was linked to limited sample diversity. Therefore, in an attempt to assess HelixMTdb sampling biases, as well as to explore this dataset more deeply, I examined in greater detail the presence or absence of HelixMTdb samples harboring each synonymous change to a third codon position (abbreviated as a ‘P3’). When considering those amino acids associated only with two-fold P3 degeneracy, or the ability to accept only two different bases at P3 without changing the protein sequence, samples diverging from the human reference sequence were present in HelixMTdb for nearly all (>97%) analyzed positions (Fig. 2a). This result indicates that the HelixMTdb does indeed cover a substantial amount of human mtDNA sequence diversity. Furthermore, our analysis demonstrates that SSNEs are unlikely to be associated with two-fold degenerate sites and that codon choice at two-fold degenerate sites governed by transitions (a purine-purine change or a pyrimidine-pyrimidine change) is unlikely to be under strong selection in humans. Since transitions at two-fold degenerate sites cause changes in local GC content, as well as slight alterations in global nucleotide content, our results also argue against substantial selection upon human mtDNA that might be based upon these factors.

**Fig. 2:**
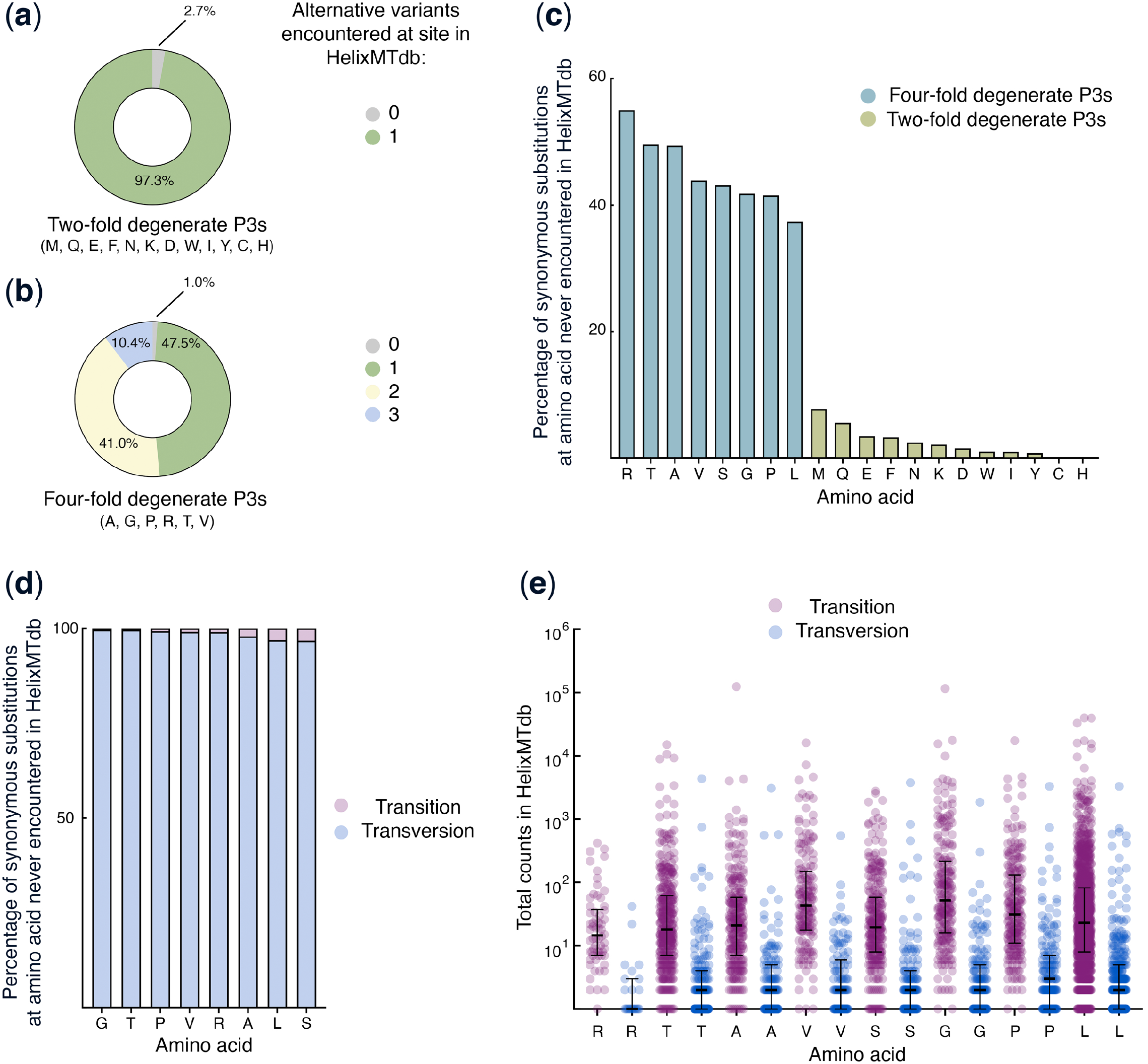
Transversion substitutions from the reference sequence are heavily depleted at four-fold degenerate third-codon positions of human mitochondrial coding sequences. **(a)** Nearly all synonymous changes from the human reference sequence can be identified within HelixMTdb at amino acid positions characterized by two-fold degenerate P3s. **(b)** Nearly all amino acid positions characterized by four-fold degeneracy at P3 could be associated with at least one base substitution in the HelixMTdb dataset. **(c)** Synonymous mutations never encountered in the HelixMTdb study are most abundant at amino acids for which at least one codon is four-fold degenerate at P3. **(d)** The vast majority of synonymous substitutions never encountered by the HelixMTdb study are transversions. **(e)** For those substitutions that are encountered in HelixMTdb, population prevalence is linked to substitution type. For each transition or transversion found in the HelixMTdb at an amino acid for which at least one codon is four-fold degenerate at P3, the HelixMTdb population count is plotted. Bar and error bars represent median of population count and interquartile range, respectively. For all comparisons of variant frequencies for a given amino acid (transition versus transversion from reference), Kolmogorov-Smirnov approximate P-values are <0.0001.

Next, I examined variation at amino acids associated only with four-fold P3 degeneracy, or accepting all nucleotides at P3 without alteration to protein sequence. Nearly all analyzed four-fold degenerate positions (99%) harbored at least one sample with a base differing from that found in the human reference (Fig. 2b), again suggesting that HelixMTdb encompasses a sizable amount of human mtDNA diversity. However, I encountered apparent limitations on base variability at some of these four-fold degenerate P3s, as all four base possibilities could be identified within HelixMTdb at only 10.4% of these mtDNA locations.

I then tested whether SSNEs might be predominantly associated with specific amino acids. I found that for the amino acids arginine, threonine, alanine, valine, serine, glycine, proline, and leucine, a more substantial fraction of synonymous changes were never seen in the HelixMTdb relative to the other amino acids (Fig. 2c). All of these amino acids can be associated with vertebrate mtDNA codons of four-fold degeneracy.

While two-fold degenerate P3s allow only transitions without altering the amino acid, four-fold degenerate P3s also permit transversions (a purine-pyrimidine change, or *vice versa*) that leave the protein code unchanged. Since amino acids with four-fold degenerate P3s were characterized by a higher number of SSNEs, I tested whether SSNEs might be more closely associated with transversions or transitions. I found that nearly all SSNEs assigned to these eight amino acids (> 96%) were linked to a potential transversion (Fig. 2d). Expanding my analysis to synonymous substitutions encountered at least once in HelixMTdb, the population frequency of a variant was also clearly linked to whether the nucleotide change at four-fold degenerate P3s was a transition or a transversion (Fig. 2e).

Two conspicuous and non-exclusive hypotheses exist regarding the notable enrichment of transversions among SSNEs at four-fold degenerate P3s. First, there may be unanticipated, yet substantial selection that acts upon P3s and leads to depletion of even synonymous transversions from the human population. Second, mutational biases related to mtDNA replication and maintenance may make transversions at degenerate P3s far less likely than transitions (Aquadro and Greenberg, 1983; Belle et al., 2005; Brown and Simpson, 1982; Kennedy et al., 2013; Kumar, 1996; Tamura and Nei, 1993; Vermulst et al., 2007; Wakeley, 1996; Zaidi et al., 2019). To address the first possibility, I further examined the extent of selection on P3s among mammals by examining the nucleotide frequencies at approximately 5 million P3s across the coding sequences of 1317 mammalian mtDNAs. Here, I also took into account the two different mitochondrial tRNAs recognizing leucine codons and the two mitochondrial tRNAs recognizing serine codons. As encountered in previous studies (Kumar, 1996; Reyes et al., 1998), guanine was depleted from mitochondrial P3s (the vast majority of which are encoded by the L-strand) for which the presence of any purine does not lead to an amino acid change (Fig. 3a), while adenine dominated at those positions. Cytosine and thymine were both well-represented at P3s for which any pyrimidine is permitted without altering the encoded amino acid. However, even considering the relative depletion of guanine from all four-fold degenerate P3s and two-fold degenerate purine P3s, guanine was nonetheless detected at thousands of P3 positions (Fig. 3b). Consequently, nucleotide frequencies at P3s appear unlikely to reflect generalized codon limitations inherent to the process of mitochondrial translation.

**Fig. 3:**
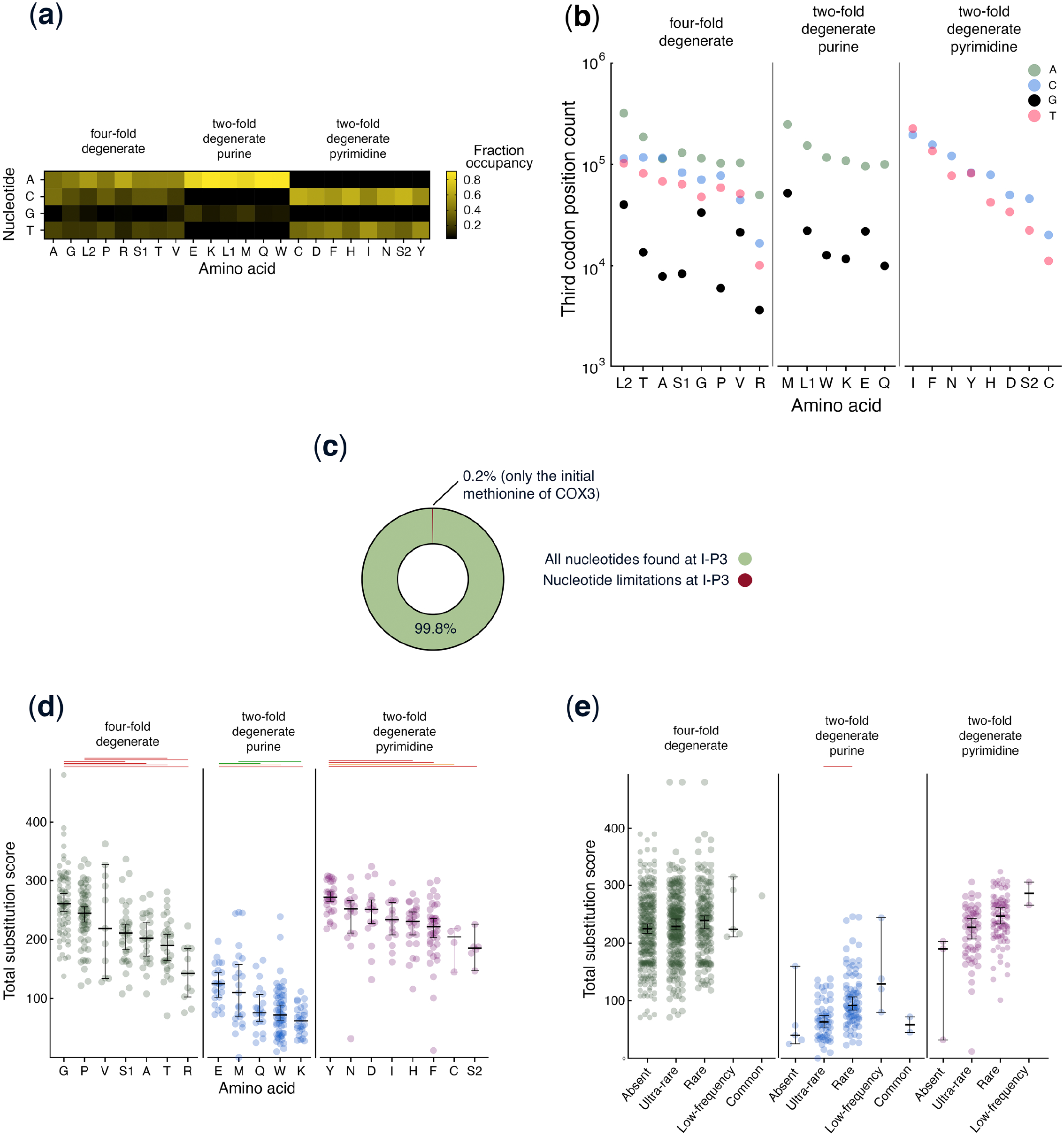
Abundant substitution and degeneracy across the third codon positions of mammals. **(a)** Base occupancy is not equally distributed among nucleotides at degenerate P3s (n ≥ 33073 instances of each amino acid are analyzed across the set of input mammalian mtDNAs). **(b)** Guanine is not excluded from recognition by any tRNA presumed to allow silent purine substitution at P3. **(c)** Nearly all I-P3s can accept all of the possible synonymous substitutions available for a particular amino acid. **(d)** Substitution is common at nearly all mammalian I-P3 positions, although the TSS distributions of I-P3s associated with amino acids can differ within each degeneracy class (four-fold degenerate, two-fold degenerate purine, or two-fold degenerate pyrimidine). Kolmogorov-Smirnov approximate P-values corrected for multiple comparisons are shown (red, ≤ 0.001; orange, ≤ 0.01; green, ≤ 0.05; no line, > 0.05). Bar and error bars represent median with 95% confidence interval. **(e)** Population frequency of synonymous variants is unlinked to TSS at four-fold degenerate I-P3s. Statistical significance is represented as in **(d)**, and the bar and error bars represent median and 95% confidence intervals.

While degeneracy at the third codon position is a general feature of mtDNA-encoded amino acids, these results are not necessarily informative about the possibility that strong selection acts upon *specific* P3s encoded by the mitochondrial genome. To further explore the extent to which individual P3s might be under selection, I focused my attention upon codons for which the first and second positions, and therefore the encoded amino acids, are 100% identical in an alignment consisting of 1251 mammals and an outlier reptile sequence (*Iguana iguana*) used to root an inferred phylogenetic tree. Leucine codons could not be included in this analysis, as degeneracy at the first codon position always led to substitution between codons recognized by the L1 and L2 tRNAs at mammalian protein alignment sites harboring only leucine. I found that 560/561 (99.8%) of the resulting set of P3s (‘I-P3s’, indicating identity of codon positions one and two throughout the alignment) can be inhabited by any nucleotide permitting synonymous substitution (Fig. 3c). Only the I-P3 of the codon annotated in humans as encoding the COX3 starting methionine appears to be totally constrained with respect to nucleotide choice, as this P3 is always occupied by guanine in mammals. Interestingly, COX3 is reported to be unique among mitochondrial polypeptides for its lack of an amino-terminal formyl-methionine (Walker et al., 2009), although whether there is a mechanistic relationship between these two observations remains to be determined. Nearly complete degeneracy among mammalian I-P3s again argues against strong selection upon synonymous nucleotide substitution within mammals based upon codon preference and the thermodynamics of base pairing, while also arguing against selection due to the presence of conserved protein binding sites.

I then calculated for each I-P3 the <u>T</u>otal <u>S</u>ubstitution <u>S</u>core (TSS), a sum of substitutions occurring at a specific site throughout an inferred mammalian phylogenetic tree (Akpinar et al., 2021). I found, with few exceptions, that nearly all four-fold degenerate, two-fold degenerate pyrimidine, and two-fold degenerate purine I-P3s have been subject to base substitution tens, or even hundreds of times, during approximately 200 million years of mammal evolution (Fig. 3d), a result quite consistent with minimal selection upon nucleotide choice at mitochondrial P3s. However, here I did note statistically significant divergence between TSS distributions at I-P3s when comparing amino acids within a given degeneracy class. These results suggest, when also considering the findings described above, that while synonymous base substitutions are not under strong selective pressure, weak selection may have helped to shape some mitochondrial third codon positions throughout the course of mammalian evolution.

Next, I asked whether a low frequency of human variants at four-fold degenerate I-P3s would correspond with lower mammal-derived TSS values of corresponding positions, a potential indicator of selection extending across mammals to humans. Here, I placed variation occurring at I-P3s into the classes ‘absent’ (zero counts among 195983 HelixMTdb samples), ‘ultra-rare’ (variant frequency < 0.01%), ‘rare’ (variant frequency < 1% and ≥ 0.01%), ‘low-frequency’ (variant frequency ≥ 1% and < than 5%) or ‘common’ (variant frequency ≥ to 5%). However, I detected no significant relationship between TSS and variant frequency for four-fold degenerate and two-fold degenerate pyrimidine I-P3s (Fig. 3e), suggesting that the prominent SSNE abundance at four-fold degenerate P3s is unlikely to be due to selection. A statistically significant link after correction for multiple testing was only observed when comparing the TSS distribution between ultra-rare and rare variants at analyzed I-P3s harboring purines, providing limited evidence of mammal-wide selection that might determine the frequency of human mtDNA variation.

Finally, I explored how the tabulation of heteroplasmic samples within the HelixMTdb might be informative regarding potential selection on synonymous and non-synonymous substitutions. Most human cells harbor hundreds of mtDNA molecules. Repeated encounters with a variant in a heteroplasmic state, where the variant is not found in all of the sequenced mtDNA molecules, is often considered to be a signal of pathogenicity, as homoplasmy of a deleterious variant is expected to lead to a fitness defect (Ye et al., 2014). However, if a synonymous change to mtDNA is neutral, whether a synonymous variant is encountered as heteroplasmic or homoplasmic should be the result of drift and a function of the number of cell divisions since the initial appearance of the novel mutation in a population (Chinnery et al., 2000; Schaack et al., 2020; Stewart and Chinnery, 2015). I plotted the frequency of heteroplasmy calls against the number of samples harboring the selected variant for those substitutions encountered in at least ten HelixMTdb samples. Synonymous variants decreased in the frequency at which they were labelled as heteroplasmic as the population frequency of encounters increased, consistent either with neutral drift toward homoplasmy or with selection (Fig. 4). However, a trend toward a higher frequency of heteroplasmy calls for non-synonymous variants than for synonymous variants was easily visualized at lower population frequencies, again consistent with a general lack of selection on human synonymous variation.

**Fig. 4:**
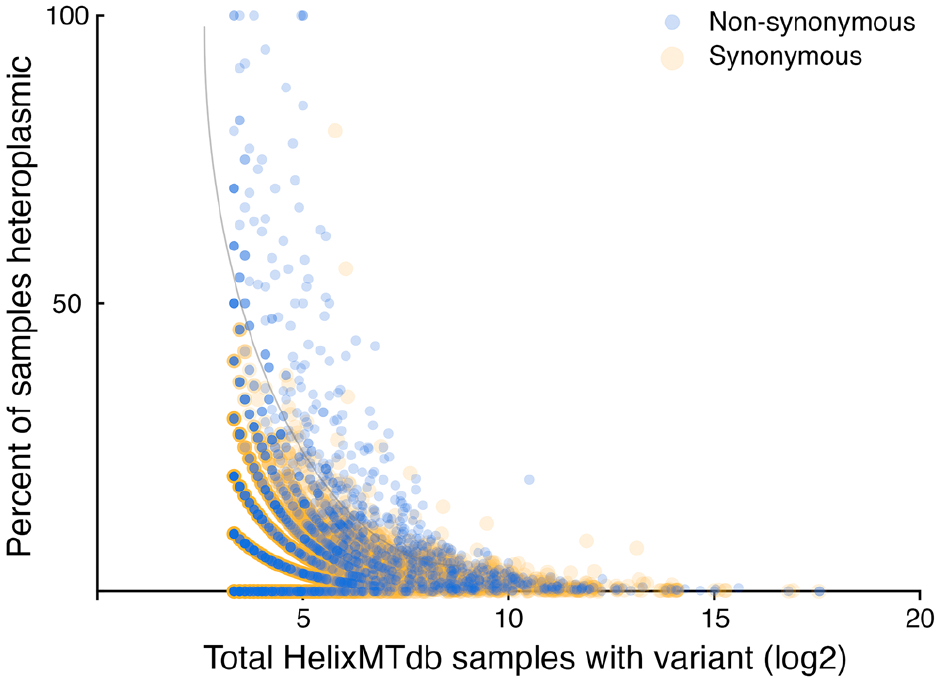
Reduced selection on synonymous substitution compared to non-synonymous substitution is indicated by an analysis of variant heteroplasmy. Only variants represented by at least 10 samples in HelixMTdb are plotted. The grey curve imposed upon the figure highlights a notable divergence in heteroplasmic sample fractions at low variant frequency that becomes apparent when comparing synonymous and non-synonymous variants.

## Conclusions and outlook

Taken together, my findings indicate that human variation at mtDNA-encoded P3s is mostly constrained by the substitution rates of each nucleotide. More specifically, variation appears highly restricted by the reduced likelihood of transversion relative to transition in mtDNA, a phenomenon well documented in earlier studies of humans and other mammals (Aquadro and Greenberg, 1983; Belle et al., 2005; Brown and Simpson, 1982; Kennedy et al., 2013; Kumar, 1996; Tamura and Nei, 1993; Vermulst et al., 2007; Wakeley, 1996; Zaidi et al., 2019). Any role for selection at synonymous sites is relatively minor, with strength of selection potentially dependent upon the specific amino acid under analysis. The low selection on codon choice encountered during my study is quite consistent with earlier, more limited analyses of mitochondrial codon choice (Castellana et al., 2011; Faith and Pollock, 2003; Jia and Higgs, 2008; Uddin and Chakraborty, 2017) and with the highly streamlined tRNA set used during mitochondrial protein synthesis.

The strong link that I have revealed between mutational biases and human variant frequencies in the large HelixMTdb dataset highlights the difficulties in assigning potential pathogenicity to non-synonymous variants based, in part, upon the variant frequency in the population (Bomba et al., 2017; Gibson, 2012; McInnes et al., 2021; Povysil et al., 2019; Zuk et al., 2014). Accordingly, attempts to link rare and *de novo* variation to disease are likely to be most successful when the mutational biases for each nucleotide can be estimated and properly taken into account.

## Data availability

The software and data that support the findings of this study are available at: https://doi.org/10.5281/zenodo.5493479.

## Methodology

### Calculation of protein changes caused by single nucleotide substitutions from the human reference sequence

The human reference mtDNA sequence and accompanying annotation (accession NC_012920.1) was downloaded from GenBank and used as input for the script ‘amino_acid_changes_caused_by_mtDNA_nucleotide_changes_in_reference.py’. This script generates a file reporting the protein sequence change associated with each simulated single base substitution to each human mitochondrial protein coding gene.

### Analysis of merged human mtDNA variation

The HelixMTdb dataset (Bolze et al., 2019) was downloaded on August 26, 2021 (https://www.helix.com/pages/mitochondrial-variant-database). The script ‘shape_HelixMTdb.py’ takes as input the HelixMTdb dataset and the output of ‘amino_acid_changes_caused_by_mtDNA_nucleotide_changes_in_reference.py’, then links HelixMTdb variation to simulated amino acid changes. A subsequent analysis of transitions and transversions from the reference sequence was performed by running the script ‘transitions_transversions.py’ on the output of ‘shape_HelixMTdb.py’.

To check the diversity of mtDNA sequences tabulated within HelixMTdb, and to further explore this dataset, we used the script ‘check_HelixMTdb_diversity.py’, which tests how many silent P3 changes from the reference sequence can be identified for each amino acid position, further classified by two-fold or four-fold P3 degeneracy. To simplify this analysis, leucine and serine codons were excluded from this analysis, since these amino acids are associated with tRNAs of both two-fold and four-fold P3 degeneracy.

### Detection of third codon position selection by analysis of mammalian mtDNAs

Records for mammalian reference mtDNAs were obtained using the Organelle Genome Resources provided by the National Center for Biotechnology Information Reference Sequence project (NCBI RefSeq, Release 207, https://www.ncbi.nlm.nih.gov/genome/organelle/) (O’Leary et al., 2016). All accessions not containing ‘NC_’ at the beginning of their accession name were removed, and this list of accessions was used to download full GenBank records [‘mammalian_mtDNA_NC_AUG_26_2021_noroot.gb’] using the NCBI Batch Entrez server (https://www.ncbi.nlm.nih.gov/sites/batchentrez). These GenBank records were analyzed by the script ‘third_codon_position_selections_all_mammals.py’ to determine total counts and frequencies of P3 bases for each amino acid. Records not containing all of the following coding sequence annotations were discarded: ‘ND1’, ‘ND2’, ‘COX1’, ‘COX2’, ‘ATP8’, ‘ATP6’, ‘COX3’, ‘ND3’, ‘ND4L’, ‘ND4’, ‘ND5’, ‘ND6’, ‘CYTB’.

To calculate TSSs (Akpinar et al., 2021) for I-P3 positions, the GenBank record for *Iguana iguana* (NC_002793.1) was added to the set of mammalian GenBank records [‘mammalian_mtDNA_NC_AUG_26_2021_Iguana_root.gb’]. The script ‘I-P3_Part_1.py’ was used to extract and align sequences from the resulting input GenBank file. Again, accessions without all of the following coding sequence annotations were discarded by this script: ‘ND1’, ‘ND2’, ‘COX1’, ‘COX2’, ‘ATP8’, ‘ATP6’, ‘COX3’, ‘ND3’, ‘ND4L’, ‘ND4’, ‘ND5’, ‘ND6’, ‘CYTB’. Accessions with any coding sequence found duplicated in another accession were also discarded. This script calls upon MAFFT v7.487 (Katoh and Standley, 2013) to align mtDNA-derived coding sequences using the FFT-NS-2 algorithm. A concatenated alignment of coding sequences output from this script was used to infer a maximum likelihood tree in RAxML-NG v1.0.3 (Kozlov et al., 2019) using a single partition, a GTR+FO+G4m model of DNA change, and a seed of 777. 10 random and 10 parsimony-based starting trees were used to initiate tree construction, and the average relative Robinson-Foulds distance (Robinson and Foulds, 1981) for the inferred trees was 0.01. 600 bootstrap replicates were generated using RAxML-NG v1.0.3, and a weighted Robinson-Foulds distance converged below a 1% cutoff value (seed of 2000). Felsenstein’s Bootstrap Proportions (Felsenstein, 1985) [‘FBP_AUG_26_2021.raxml.support] and the Transfer Bootstrap Expectations (Lemoine et al., 2018) [‘TBE_AUG_26_2021.raxml.support’] were calculated and used to label the best scoring tree [‘T3_AUG_26_P3_REV.raxml.bestTree’]. This tree was used for downstream analyses after using FigTree 1.4.4 (https://github.com/rambaut/figtree/releases) to place the root upon the branch leading to *Iguana iguana* [‘AUG_26_P3_REV_tree_rooted_Iguana_iguana.nwk’].

This rooted tree and the coding sequence alignments for each mitochondria-encoded protein were used to determine the TSSs associated with each class of P3 and to determine which bases can occupy mammalian I-P3 sites. Most four-fold degenerate P3s within codons with identical first and second positions were analyzed using the ‘I-P3_Part_2_AGPRTV.py’ script, most two-fold degenerate purine P3s were analyzed with ‘I-P3_Part_2_EKMQW.py’, and most two-fold degenerate pyrimidine P3s were analyzed with ‘I-P3_Part_2_CDFHINY.py’. Any amino acids positions encoded by tRNA L1 in all mammals and the outgroup, and therefore harboring T and T at the first and second codon positions in all samples, were sought by script ‘I-P3_Part_2_L1.py’. Similarly, amino acids positions encoded by tRNA L2 in all mammals and the outgroup, and therefore harboring C and T at the first and second codon positions in all samples, were would have been identified and analyzed by script ‘I-P3_Part_2_L2.py’. The two-fold degenerate P3s associated with the use of tRNA S2 were analyzed by script ‘I-P3_Part_2_S2.py’, and the four-fold degenerate P3 associated with serines encoded by tRNA S1 were analyzed by script ‘I-P3_Part_2_S1.py’. Within each of these scripts, MAFFT v7.487 was used to perform alignments using the FFT-NS-2 algorithm, script ‘ungap_on_reference.py’ v1.0 (Dunn, 2021) was used to ungap alignments based on the human reference sequence, ancestral character predictions were made at internal nodes using RAxML-NG v1.0.3 (Kozlov et al., 2019), and seqkit v0.16.1 (Shen et al., 2016) was used while formatting the node names associated with ancestral sequences.

### Comparison of total substitution scores to HelixMTdb dataset

Output of the above-mentioned scripts was combined into new tables and further annotated based upon whether the P3 data were obtained from two-fold degenerate purine sites [‘two_fold_AG_P3_ALL.csv’], two-fold degenerate pyrimidine sites [‘two_fold_CT_P3_ALL.csv’], or four-fold degenerate sites [‘four_fold_P3_ALL.csv’]. Further processing to quantify any relationship between TSS and general P3 type, TSS and amino acid, the link between (non-)synonymous mutation and sample counts in HelixMTdb was performed using the script ‘I-P3_Part_3.py’.

### Statistical analyses

The Kolmogorov-Smirnov test, a non-parametric approach testing the hypothesis that two samples are drawn from different populations, was used to compare the total variant counts of synonymous and non-synonymous substitutions, to contrast total transversion versus transition counts for specific amino acids, and to compare TSS distributions between amino acids or frequency classes. Kolmogorov-Smirnov analyses were performed in Prism 9.2.0 or SciPy v1.7.1 (Virtanen et al., 2020). When analyzing I-P3 TSS values among different frequency classes, TSS distributions are arranged by amino acid, which are then compared by Kolmogorov-Smirnov testing only within a given category (four-fold degenerate, two-fold degenerate purine, two-fold degenerate pyrimidine). Moreover, the ‘common’ frequency class was not subject to statistical testing due to the limited number of variants found within this category. Correction for multiple testing (Bonferroni correction) was accomplished by multiplying each single test P-value by the number of tests.

## Acknowledgements

Funding for this project was obtained from the Sigrid Jusélius Foundation (Senior Researcher Grant to C.D.D.) and the European Research Council (ERC Starting Grant RevMito 637649 to C.D.D.). I appreciate the computational support provided by the Center for Scientific Computing, Finland (Puhti supercomputer). I thank Ani Akpinar for assistance with HelixMTdb processing and for critical comments on the analysis. I also thank Gülayşe İnce Dunn and Svetlana Konovalova for helpful manuscript comments.

## Conflict of Interests

The authors declare that they have no conflict of interest.

## Notes

https://doi.org/10.5281/zenodo.5493479

### Competing Interest Statement

The authors have declared no competing interest.

### Summary of Updates

Changes to the text, along with additional analyses, are found in this new version of the manuscript. Peer review comments and responses associated with this revised version were posted on bioRxiv by Review Commons on September 8, 2021.

